# The β2-adrenergic receptor (ADRB2) entrains circadian gene oscillation and diurnal responses to virus infection in CD8^+^ T cells

**DOI:** 10.1101/2024.03.12.584692

**Authors:** Drashya Sharma, Kira A. Kohlbach, Robert Maples, J. David Farrar

## Abstract

Adaptive immune cells are regulated by circadian rhythms (CR) under both steady state conditions and during responses to infection. Cytolytic CD8^+^ T cells display variable responses to infection depending upon the time of day of exposure. However, the neuronal signals that entrain these cyclic behaviors remain unknown. Immune cells express a variety of neurotransmitter receptors including nicotinic, glucocorticoid, and adrenergic receptors. Here, we demonstrate that the β2-adrenergic receptor (ADRB2) regulates the periodic oscillation of select core clock genes, such as *Per2* and *Bmal1*, and selective loss of the *Adrb2* gene dramatically perturbs the normal diurnal oscillation of clock gene expression in CD8^+^ T cells. Consequently, their circadian-regulated anti-viral response is dysregulated, and the diurnal development of CD8^+^ T cells into variegated populations of cytolytic T cell (CTL) effectors is dramatically altered in the absence of ADRB2 signaling. Thus, the *Adrb2* directly entrains core clock gene oscillation and regulates CR-dependent T cell responses to virus infection as a function of time-of-day of pathogen exposure.

**One Sentence Summary:** The β2-adrenergic receptor regulates circadian gene oscillation and downstream daily timing of cytolytic T cell responses to virus infection.

## 1. Introduction

The immune system is tightly regulated by a variety of signals that control responses to foreign pathogens while suppressing autoreactivity. In addition to the well characterized immune regulators such as antigen receptors and cytokines, circadian rhythms (CRs) also play a role in modulating immune responses as a function of light/dark cycles (1–4). However, the molecular pathways that control time-of-day dependent immune responses are not well understood.

In higher vertebrates, light sensing is a powerful entrainment signal for the central clock within the brain (5). However, time regulation of organ systems outside the brain requires signals delivered to them from the brain’s central clock, i.e., the suprachiasmatic nucleus (SCN). These signals come in the form of soluble neurotransmitters that are released in a diurnal fashion to regulate peripheral clocks (6). While the oscillatory pattern of core clock gene expression is directly regulated by light inputs within the SCN, peripheral tissue clock oscillation relies upon local and systemic neurotransmitters to provide circadian timing of expression. There is some evidence to suggest that immune cell functions are regulated by systemic neurotransmitters. For example, lymphocyte recirculation has been shown to follow a 24 hr periodic ebb and flow through secondary lymphatic tissues, and this trafficking was controlled by both norepinephrine (NE) and glucocorticoids (GCs) (7, 8). Indeed, a variety of neurotransmitters have been shown to regulate various aspects of immune cell function, including GCs, NE, and acetycholine (Ach), all of which are generally immunosuppressive in nature (9–15). These neurotransmitters are released in a diurnal fashion to entrain the periodic oscillation of a myriad of global biological functions including immune cell regulation (8, 16, 17). Furthermore, the core clock genes, such as Bmal1 and Per/Cry oscillate in virtually all immune cells that have been examined (18–21). However, the signaling pathways that entrain core clock gene oscillation and downstream functionality in immune cells has not been described.

Our recent study measuring transcriptional responses of T cells to viral infection uncovered a central role of the β2-adrenergic receptor (*Adrb2*) in regulating multiple cytokine- and T cell effector-pathway genes (22). In addition, we observed some unexpected pathways to be differentially regulated in *Adrb2* knockout CD8 T cells such as serotonin responsiveness, prion diseases, and circadian rhythms, which underscore the many aspects of neural regulation of immune function. In this study, we focused on the role of the *Adrb2* in regulating downstream circadian rhythms, and we uncover an unprecedented role for the *Adrb2* in directly entraining circadian gene oscillation as a function of light/dark cycles. The oscillatory nature of CR gene expression correlated with time-of-day developmental variegation of responding T cells as they differentiated in response to virus infection, which was markedly disrupted in the absence of the *Adrb2*. This study represents the first demonstration of a direct entrainment pathway for CR gene oscillation in immune cells.

## 2. Results

### 2.1 Deletion of the *Adrb2* alters CR gene expression in CD8^+^ T cells responding to virus infection in vivo

CD8^+^ T cells express the ADRB2, and we and others found that signaling through this receptor by NE or other β-agonists suppresses cytokine secretion and lytic activity in vitro and in vivo (13, 23, 24). However, the role of adrenergic signaling in regulating peripheral T cell development during infections has not been examined until recently. To this end, our recent study demonstrated a key role for the *Adrb2* in regulating downstream transcriptional pathways engaged in CD8^+^ T cells as they divide and respond to endogenous virus infection (22). This study was the first to reveal unique pathways regulated by adrenergic signaling in T cells, and in the current study, we focused our analysis on the circadian rhythm pathway. In this experiment, published in Estrada et. al. (22), purified CD8^+^ Clone4 (HA-specific TCR-Tg T cells) wildtype and Clone4 x *Adrb2*^-/-^ T cells were co-transferred to WT Balb/c recipients. The next day, animals were injected with either PBS or infected with a sub-lethal dose of VSV-HA. Cells were purified from each of 3 recipient animals at each time point, and RNASeq analysis was performed on highly purified Clone4 CD8^+^ T cells (Primary data available in GEO at accession number GSE102478).

In our previous report (22), EdgeR analysis (25) identified over 320 genes that were differentially expressed between WT and *Adrb2*^-/-^ cells at any of the timepoints, including day 0. A comprehensive integrative pathway analysis of these constituent genes identified a variety of known effector and cytokine-regulated pathways differentially regulated in *Adrb2*^-/-^ cells. One pathway that stood out was the KEGG CR pathway, which was significantly regulated at multiple timepoints of infection. From that dataset, an expression heatmap of known core CR genes is shown in Fig. 1A. We observed significant differential expression of the key circadian genes *Arntl2* (*Bmal2*, Fig. 2D), *Cry1* (Fig. 2A), *Per2* (Fig. 1C), *Per3* (Fig. 2F), and *Fbxl3* (Fig. 1C) at various timepoints of infection (statistical analysis presented in Supplemental Fig.1). In contrast, other circadian genes such as *Clock* and *RORa* were not differentially expressed at any timepoint (Figs. 1B and 2G, respectively). This experiment was a co-transfer protocol, where both WT and *Adrb2*^-/-^ responded to the virus infection simultaneously within the same WT hosts. Although the cells were harvested at different days post-infection, collection and purification of the cells was performed at approximately 2hrs into the light cycle (ZT+2, by convention) on each day. Thus, these data suggest that the *Adrb2* regulates expression of some core CR genes in virus-specific CD8^+^ T cells both at baseline and at incremental times after infection.

**Figure 1.**
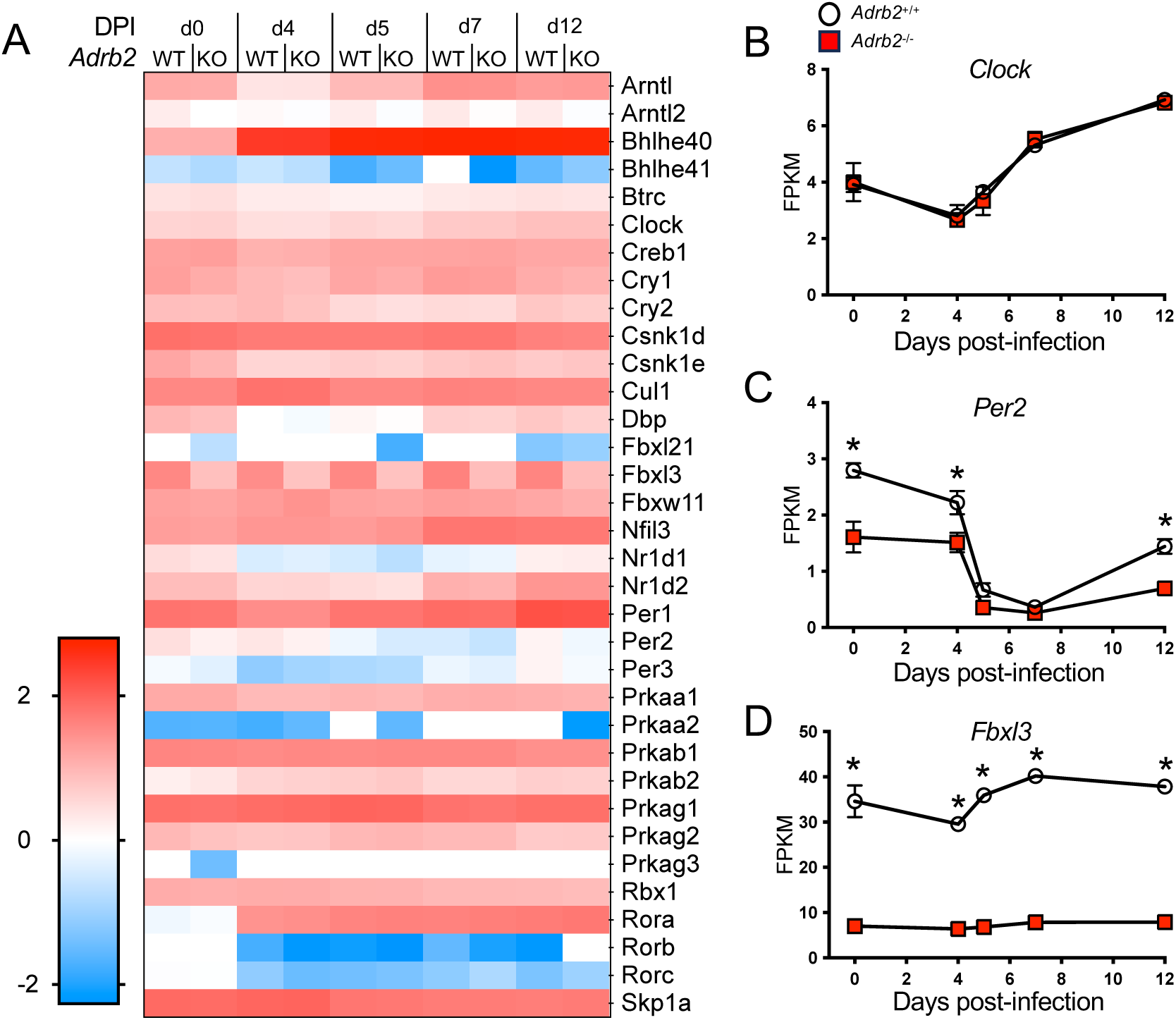
Differential core clock gene expression in *Adrb2*^-/-^ CD8^+^ T cells in response to virus infection. Data are derived from Estrada et. al.(22) comparing gene expression values (FPKM) from adoptively transferred WT and *Adrb2*^-/-^ CD8^+^ T cells purified from VSV-infected hosts. (**A**) Heatmap analysis of KEGG-annotated circadian rhythm pathway genes expressed in CD8^+^ T cells at the indicated days post infection. (**B**-**D**) Expression of *Clock*, *Per2*, and *Fbxl3* in WT vs *Adrb2*^-/-^ CD8^+^ T cells at the indicated days post infection. *p<0.05 by EdgeR analysis.

**Figure 2.**
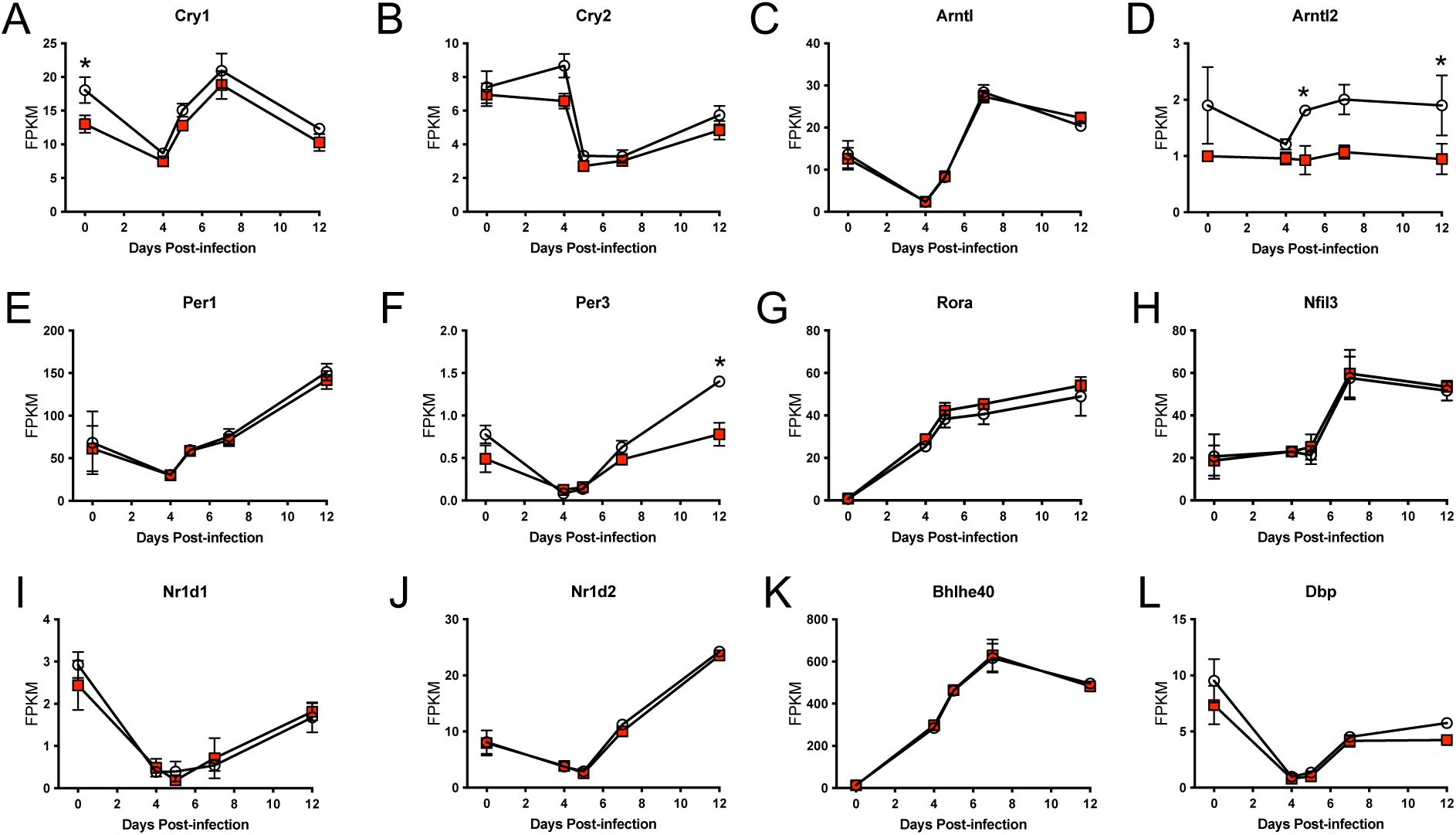
Differential expression of select core clock genes in *Adrb2*^-/-^ CD8^+^ T cells in response to virus infection. Data are derived from Estrada et. al.(22) comparing gene expression values (FPKM) from adoptively transferred WT (open circles) and *Adrb2*^-/-^ (red squares) CD8^+^ T cells purified from VSV-infected hosts. Gene expression values of (A) Cry1, (B) Cry2, (C) Arntl, (D) Arntl2, (E) Per1, (F) Per3, (G) Rora, (H) Nfil3, (I) Nr1d1, (J) Nr1d2, (K) Bhlhe40, and (L) DBP are displayed at the indicated times post-infection. *p<0.05 by EdgeR analysis.

In addition to a marked dysregulation of a subset of CR genes in *Adrb2*^-/-^ cells, we also observed that many of the core CR genes were differentially expressed in CD8^+^ T cells on incremental days following infection including all of the cryptochrome genes (*Per1-3* and *Cry1/2*), as well as *Arntl*, *Rora*, *Nfil3*, *Nr1d1*, *Nr1d2*, *Bhlhe40*, and *Dbp* (Fig. 2). For example, *Bmal1* (*Arntl*) was significantly suppressed at d4 post-infection and dramatically increased 100-fold by d7 (Fig. 2C). In contrast, *Rora* steadily increased through d12 compared to baseline levels prior to infection. These data indicate an unexpected regulation of core CR genes in CD8^+^ T cells as they divide and differentiate in response to virus infection. Given the dysregulation of some of these genes in the absence of the *Adrb2*, we wished to determine if the *Adrb2* directly regulated their oscillation through a 24 hour light/dark cycle.

### 2.2 The *Adrb2* controls diurnal oscillation of core CR genes and other immune related genes in CD8^+^ T cells

To create a more physiological model for the studies described below, we crossed the *Adrb2-fx* mouse (on C57Bl/6 background) (26) to the CD8α-Cre (27), which expresses Cre late in CD8^+^ T cell development in the thymus (27). This model selectively deletes floxed targets (such as the *Adrb2*) in peripheral CD8^+^ T cells, preserving ADRB2 signaling in other cell types and tissues. Thus, the upstream oscillation of NE and downstream response to NE remains intact in all cells except CD8^+^ T cells. To determine if the *Adrb2* directly regulated core clock gene expression in CD8^+^ T cells, we bred the *Adrb2*-fx x CD8α-Cre (<u>now referred to as</u> *<u>Ad-fx-Cre</u>*) to the Per2::Luc mouse, which reports on PER2 protein within cells and tissues (28), allowing for the measurement of PER2 oscillation.

To assess the relative temporal expression of PER2 protein, cohorts of WT Per2::Luc and *Ad-fx-Cre* x Per2::Luc mice were acclimated for 3-5 weeks to 12:12 light/dark cycles in circadian cabinets that were staged 6 hrs apart. By convention, ZT0 initiates the beginning of the light cycle. After acclimation, animals were sacrificed at ZT+4, +10, +16, and +20 corresponding to their individual cabinet time zone. Single cell suspensions were prepared from spleen, and luciferase activity was measured from purified CD8^+^ T cells. We observed significant PER2 protein oscillation in WT CD8^+^ T cells that was significantly phase shifted in *Adrb2-fx-Cre* cells (Fig. 3A). In addition to PER2 protein, total RNA was isolated from purified CD8^+^ T cells, and core CR and other select transcription factor gene expression was quantified by qPCR. As expected, *Clock* was uniformly expressed at all timepoints, and was not affected by deletion of the *Adrb2* (Fig. 3E, statistical analyses presented in Supplemental Table 1). However, in contrast to the effects on PER2 protein oscillation, we found that ablation of *Adrb2* completely disrupted significant oscillation of *Per2* and *Per3* mRNA with no impact on *Per*1 (Fig. 3B). Likewise, *Bmal2* (*Arntl2*) rhythmicity was also abolished in *Adrb2-fx-Cre* cells (Fig. 3C, right panel). *Cry1* and 2 gene oscillation amplitude was modest, even in WT cells, with only *Cry2* rhythmicity disrupted in the absence of the *Adrb2* (Fig. 3D). Thus, these data suggest that oscillation of the core cryptochrome *Per2* and *3* genes are selectively controlled by ADRB2 signaling in CD8^+^ T cells.

**Figure 3.**
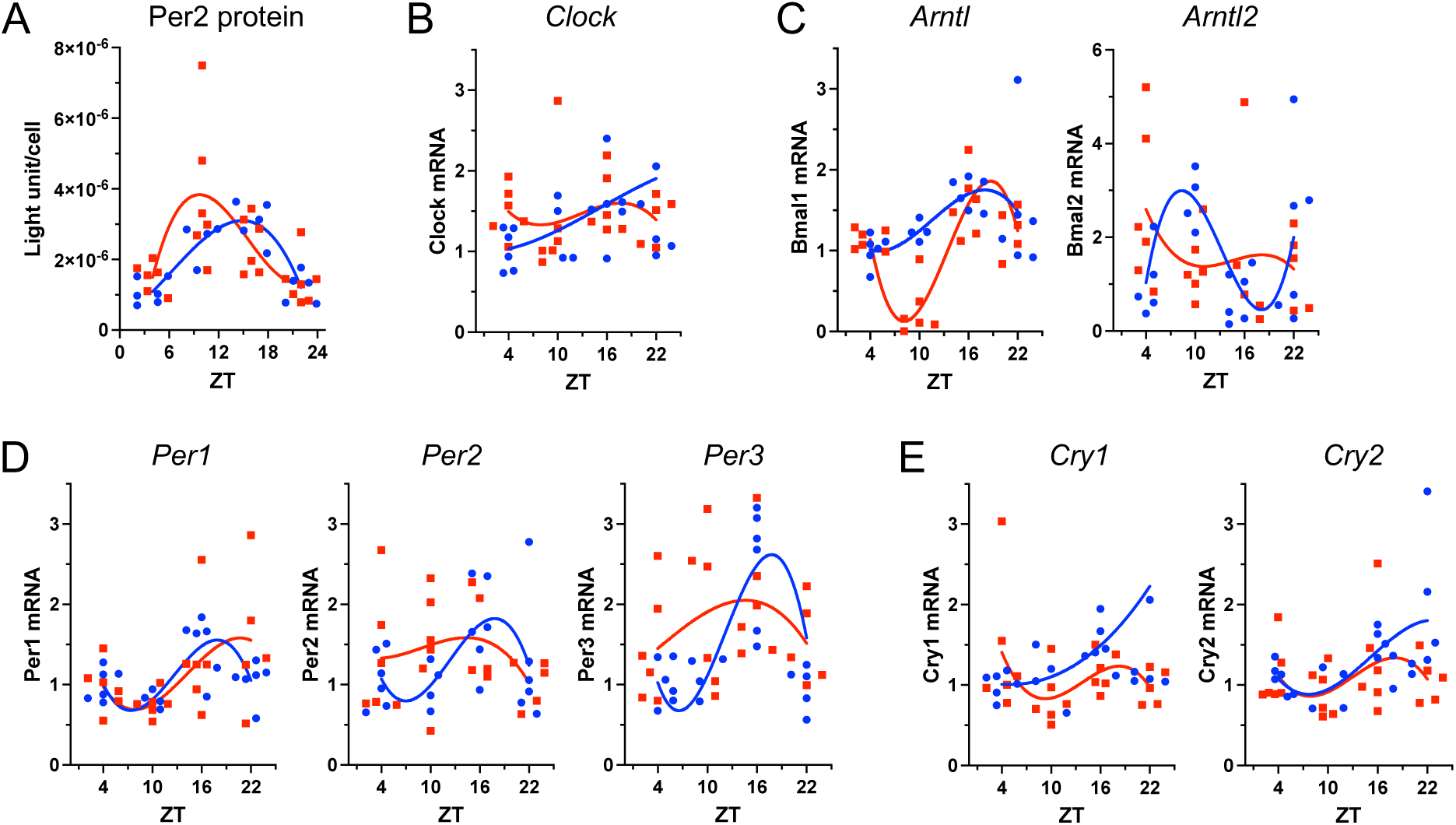
The *Adrb2* regulates periodic core clock gene oscillation in CD8^+^ T cells. 8-10 week old WT (blue circles) and *Ad-fx-Cre* x Per2::Luc (red squares) mice were acclimated for 3 weeks in light chambers to a standard 12:12 light/dark cycle staged 6 hrs apart. CD8^+^ T cells were isolated from spleens at ZT +4, +10, +16, and +22. (A) Per2 protein luciferase activity was measured by standard luminescence assay. (B-E) Total RNA was extracted from CD8^+^ T cells, and specific gene expression for *Clock* (B), *Arntl* (*Bmal*) (C), *Per* (D), and *Cry* (E) genes were quantified by qPCR. Each symbol represents a separate animal (n=4-6), and statistical significance of rhythmicity was determined by Circacompare. Oscillations were considered rhythmic at p<0.05, and evaluations of rhythmicity are indicated above each graph. If oscillations in both WT and KO cells were rhythmic, then differences in mesor, phase and amplitude were calculated and considered significantly different at p<0.05. Statistical analysis data are presented in Supplemental Table 1.

**Table 1:**
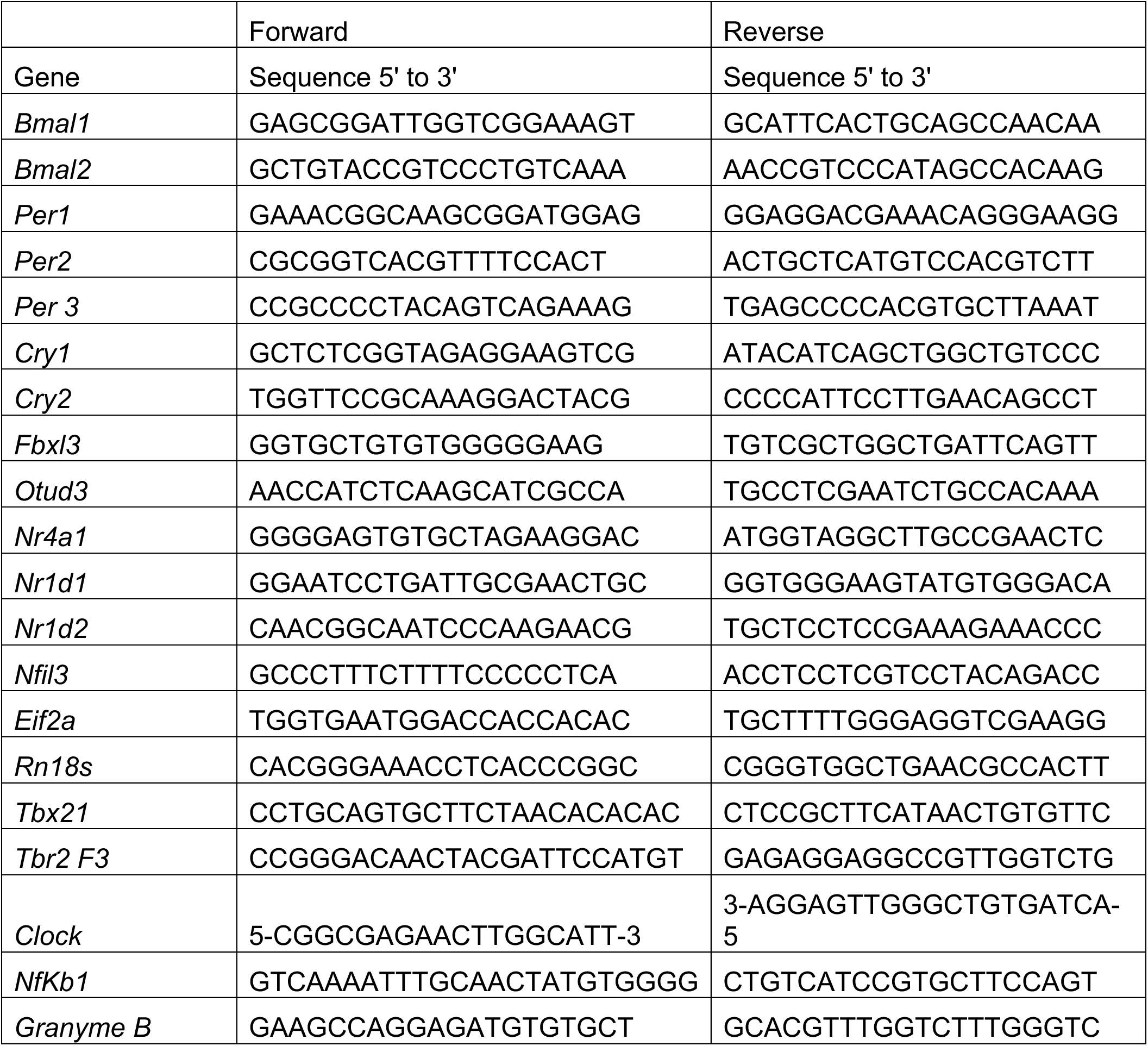
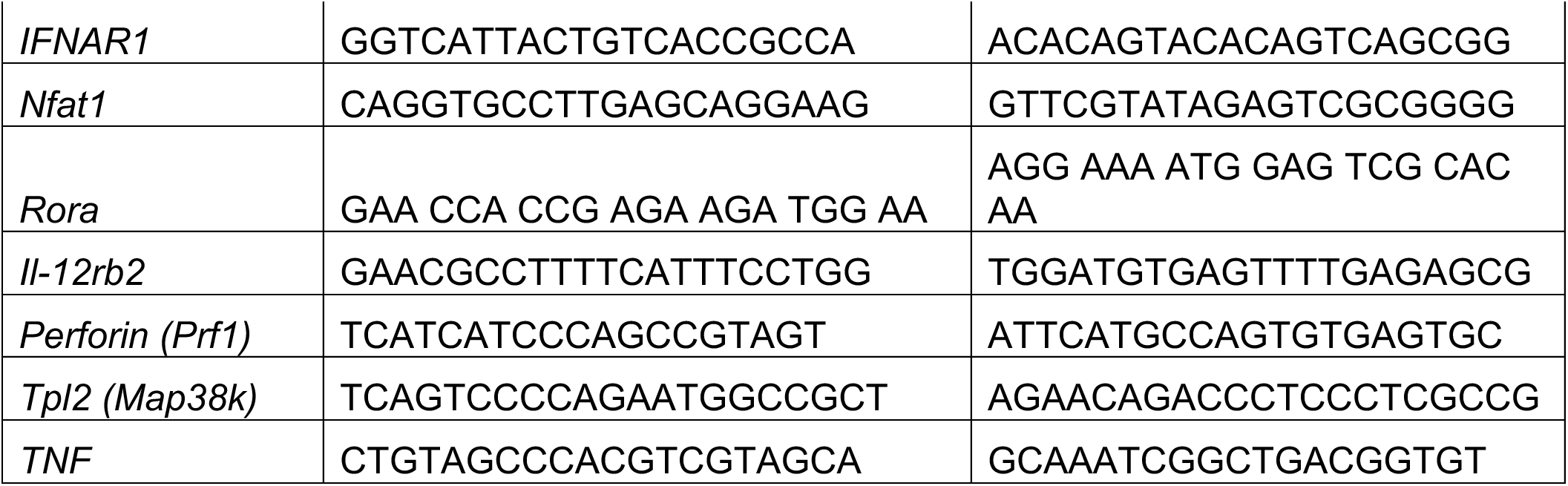
Primer sequences used in qPCR.

In measuring other core CR genes, we were surprised to find that some genes adopted alternative oscillation phases in the absence of the *Adrb*2. For example, *Bmal1* (*Arntl*) and *Nr4a1* were significantly rhythmic in both WT and KO cells but differed significantly in mesor (Fig. 3C left panel and Fig. 4A). Likewise, *Fbxl3* adopted a completely alternate phase and amplitude in *Adrb2-fx-Cre* compared to WT CD8^+^ T cells (Fig. 4A). In contrast, other core CR genes, such as *Rora*, *Nfil3*, *Nr1d1* and *Nr1d2* were arrhythmic in WT cells but adopted a unique rhythmicity in *Adrb2-fx-Cre* cells (Fig. 4A). In addition to core CR genes, we measured potential downstream genes involved in T cell activation and function. We observed similar disruptions and alterations in rhythmicity in some of these genes, such as, *Nfkb1*, *Tpl2*, and cytokine/effector genes *Gzmb*, *Pfn*, *Ifng*, and *Tnfa*. Of note, the main effector/memory T cell transcription factors *Tbet* (*Tbx21*) and *Eomes* (*Tbr2*) were significantly rhythmic in WT cells while arrhythmic in *Adrb2-fx-Cre* (Supplemental Table 1), although their overall amplitude throughout the 24 hr period was approximately 2-fold from the mesor. Collectively, we conclude from these data that intrinsic expression of the *Adrb2* is required for the rhythmic oscillation of key CR genes and some downstream T cell effector genes. In the absence of the *Adrb2*, these rhythmic patterns are either disrupted, altered, or abolished altogether, depending upon the specific gene. Thus, the ADRB2 is a critical intrinsic regulator of CR gene rhythmicity in CD8^+^ T cells.

**Figure 4.**
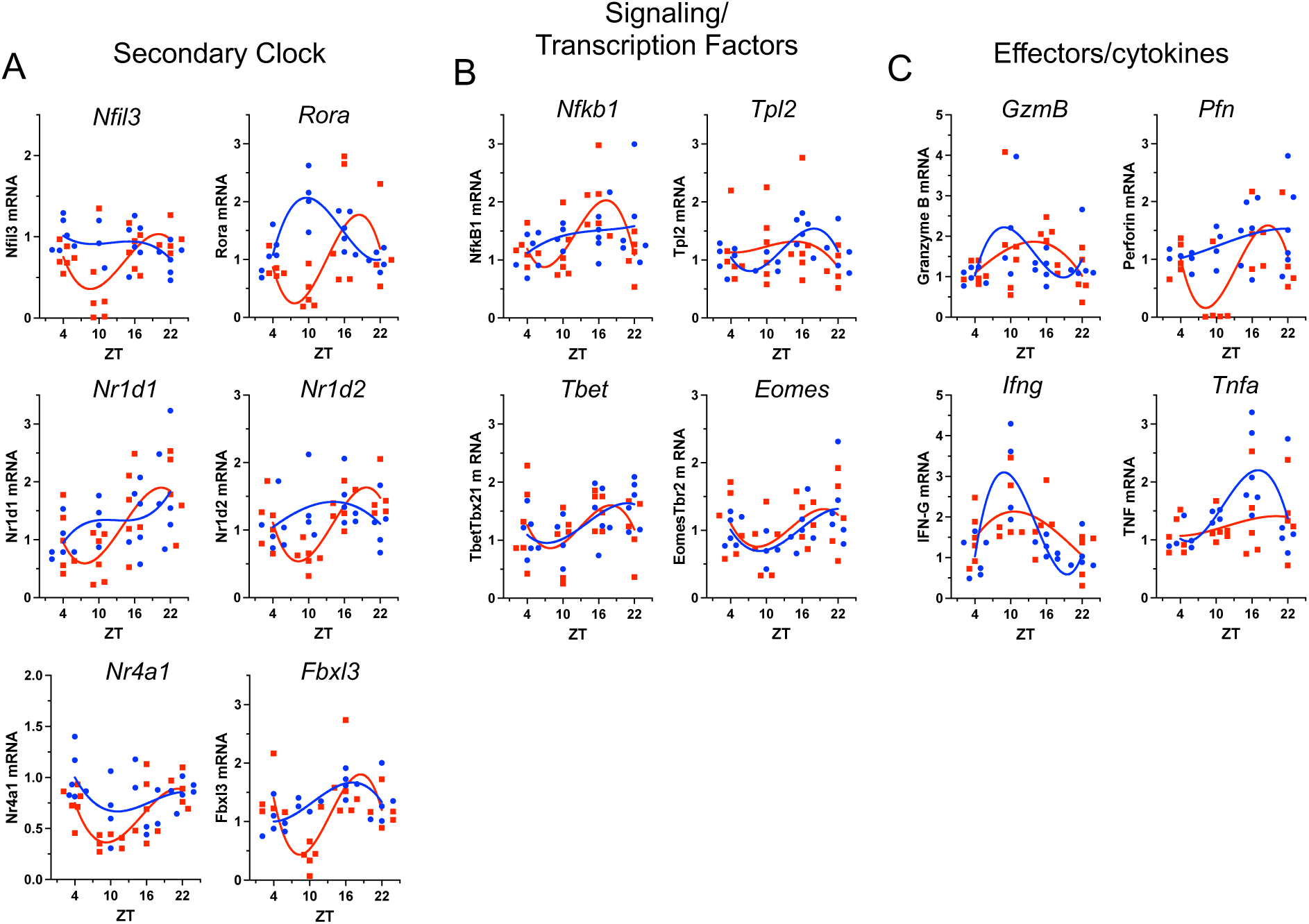
The *Adrb2* regulates periodic clock-controlled gene oscillation in CD8^+^ T cells. WT (blue circles) and *Ad-fx-Cre* x Per2::Luc (red squares) CD8^+^ T cells were isolated from spleens at ZT +4, +10, +16, and +22 as described in Fig. 2. Total RNA was extracted from CD8^+^ T cells, and mRNA transcripts for select genes were quantified by qPCR. (A) Secondary core clock genes Nfil3, *Clock* (B), *Arntl* (*Bmal*) (C), *Per* (D), and *Cry* (E) genes were quantified by qPCR. Each symbol represents a separate animal (n=4-6), and statistical significance of rhythmicity was determined by Circacompare. Oscillations were considered rhythmic at p<0.05, and evaluations of rhythmicity are indicated above each graph. If oscillations in both WT and KO cells were rhythmic, then differences in mesor, phase and amplitude were calculated and considered significantly different at p<0.05. Statistical analyses of data are presented in Supplemental Table 1.

### 2.3 The *Adrb2* regulates cytokine secretion potential and CD8^+^ T cell subset variegation as a function of time-of-day of virus infection

If the *Adrb2* regulates CD8^+^ T cell intrinsic clock gene oscillation as we propose, does this correlate with their functional responses to viral infection? To address this, we compared the effector T cell response to VSV infection between WT and Ad-fx-Cre mice. Here, cohorts of WT and Ad-fx-Cre mice were infected with VSV-OVA at ZT 0, 6, 12, and 18 within a 12:12 light/dark cycle. The VSV-OVA recombinant virus expresses chicken ovalbumin used as a traceable T cell antigen (Ag). All animals were harvested at the same time of day on d7 after infection, and splenocytes were stimulated in vitro with PMA + ionomycin to elicit cytokine expression (29, 30). Stimulated cells were then stained with a panel of 28 metal-conjugated antibodies that included a labeled MHC class I MHC-I H2-K^b^ tetramer loaded with the OVA 257-264 peptide. This tetramer is loaded with the OVA peptide that is expressed by the VSV-OVA virus, and it specifically binds only CD8^+^ T cells that have specificity for the virus (31, 32). Cells were then measured by high-dimensional CyTOF, and data were analyzed on the OMIQ data analysis platform (app.omiq.ai, Fig. 5). As expected, we identified a population of CD8^+^/Tetramer^+^ cells that developed specifically in response to infection (Fig. 3A and B, blue gate). We then quantified the percentages of cytokine secreting cells within the Tetramer^+^ population from infected animals. We measured IL-2, IL-6, IL-10, IL-17A, IFN-γ, TNF-α, along with Perforin and Granzyme B, and an example gating is shown for the expression of Granzyme B in Fig. 3C. The cytokines IL-6, IL-10, and the lytic enzymes Perforin and Granzyme B (Fig. 5D) displayed no significant rhythmicity of expression, nor were there significant differences in their expression as a function of time-of-day of virus exposure. In stark contrast, IL-2 expression was unique among this group in that it displayed dramatic rhythmicity, which was not affected by the loss of the *Adrb2* (Fig. 5E). However, the rest of the cytokines that we analyzed (IFN-g, TNF-a, and IL-17A) exhibited time-of day dependent rhythmicity that was lost in the *Ad-fx-Cre* cells, and an example of this pattern is shown for IFN-γ^+^ and TNF-α^+^ cells in Fig. 5F and G. These data demonstrate a role for the *Adrb2* in selectively regulating time-of-day dependent CD8^+^ T cell cytokine responses to viral infection.

**Figure 5.**
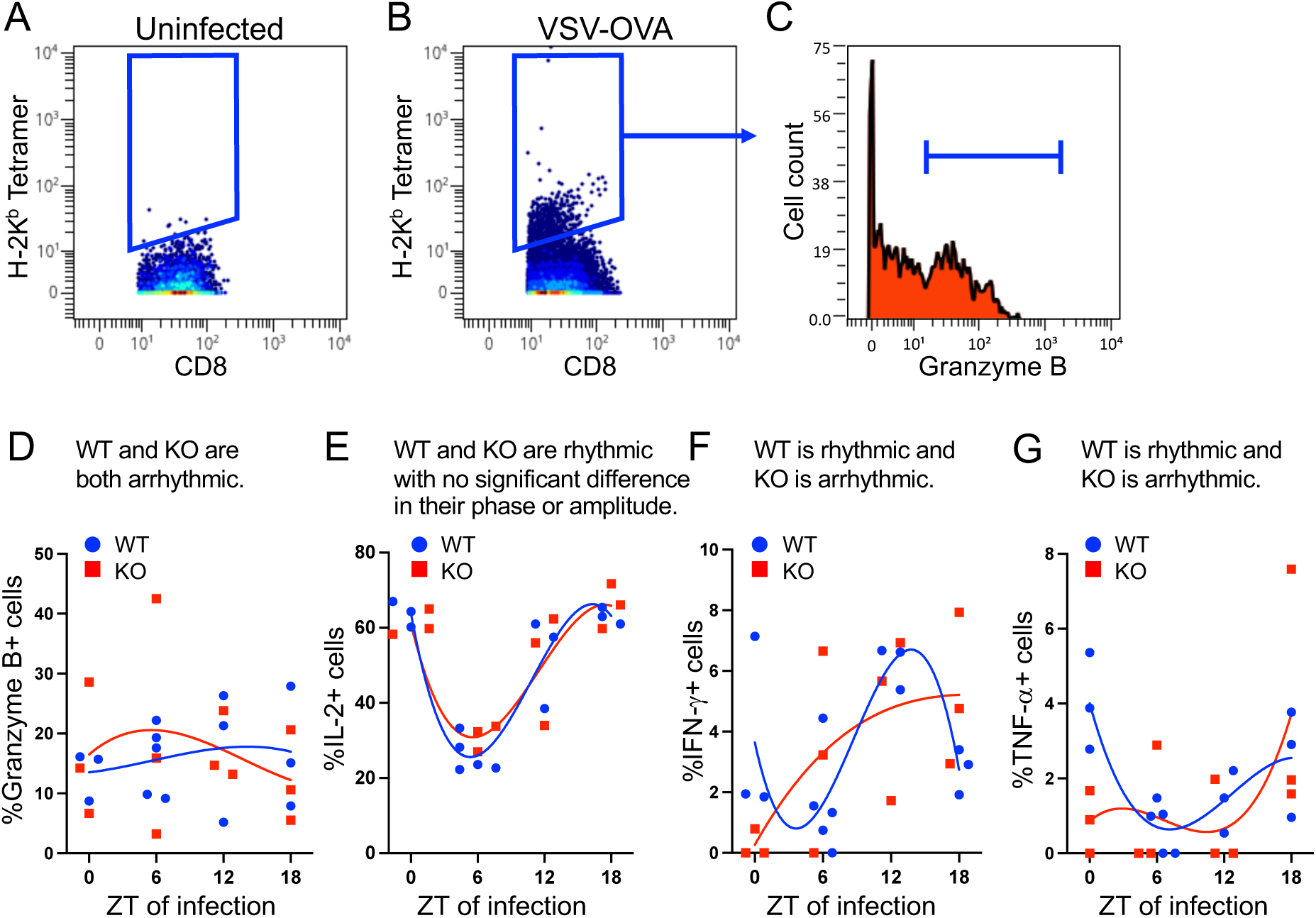
The *Adrb2* controls the rhythmicity of select cytokine secretion by virus-specific CD8^+^ effector cells. WT and Ad-fx-Cre cohorts (n=3-5/group) were infected with VSV-OVA at the indicated ZT times and allowed to recover to d7. All animals were harvested at the same time on d7, and CyTOF analysis was performed on in vitro stimulated splenocytes. (A and B) Data were gated on CD8 and K^b^ tetramer positive cells (blue gate) from uninfected (A) and VSV-OVA infected (B) animals, and the data were further gated on cytokine- or effector molecule-secreting cells (blue bar gate), e.g. Granzyme B (C). CD8^+^/Tetramer^+^ cells from infected animals were quantified for the percentage of Granzyme B^+^ (D), IL-2^+^ (E), IFN-γ^+^ (F), and TNF-α^+^ (F) cells, and each symbol represents the average of replicate analysis from individual animals. Statistical significance of rhythmicity was determined by Circacompare and is described above each graph.

We wished to correlate the rhythmic development of cytokine-secreting cells observed in Fig. 5 with the expansion of Ag-specific cells. Here, cohorts of WT and *Ad-fx-Cre* mice were acclimated and infected with VSV-OVA at the ZT times indicated in Fig. 5. Spleens were harvested at d7 at the same time of day, and cells were stained with fluorochrome-conjugated antibodies to detect CD4 and CD8, and a fluorchrome-conjugated Kb-OVA-Tetramer to identify virus-specific CD8^+^ T cells. First, in assessing bulk T cells, we found no significant oscillation in either total CD4^+^ or CD8^+^ cells as a function of time of virus infections (Fig. 6A-B). The expansion of antigen-specific cells in the infected cohort was robust (Fig. 6C), yet we also found no significant differences in either the percentages (data not shown) or total numbers (Fig. 6D) of Ag-specific cells between WT and *Adrb2*-fx mice when infected at different times of day. Thus, the expansion of total Ag-specific cells was not regulated by the time of day at which the animals were exposed to virus. Further, the differences in numbers of IFN-γ-and other cytokine-secreting effector cells as a function of time of infection seen in Fig. 5 cannot be accounted for by different numbers of Ag-specific cells which expand over the course of the infection.

**Figure 6.**
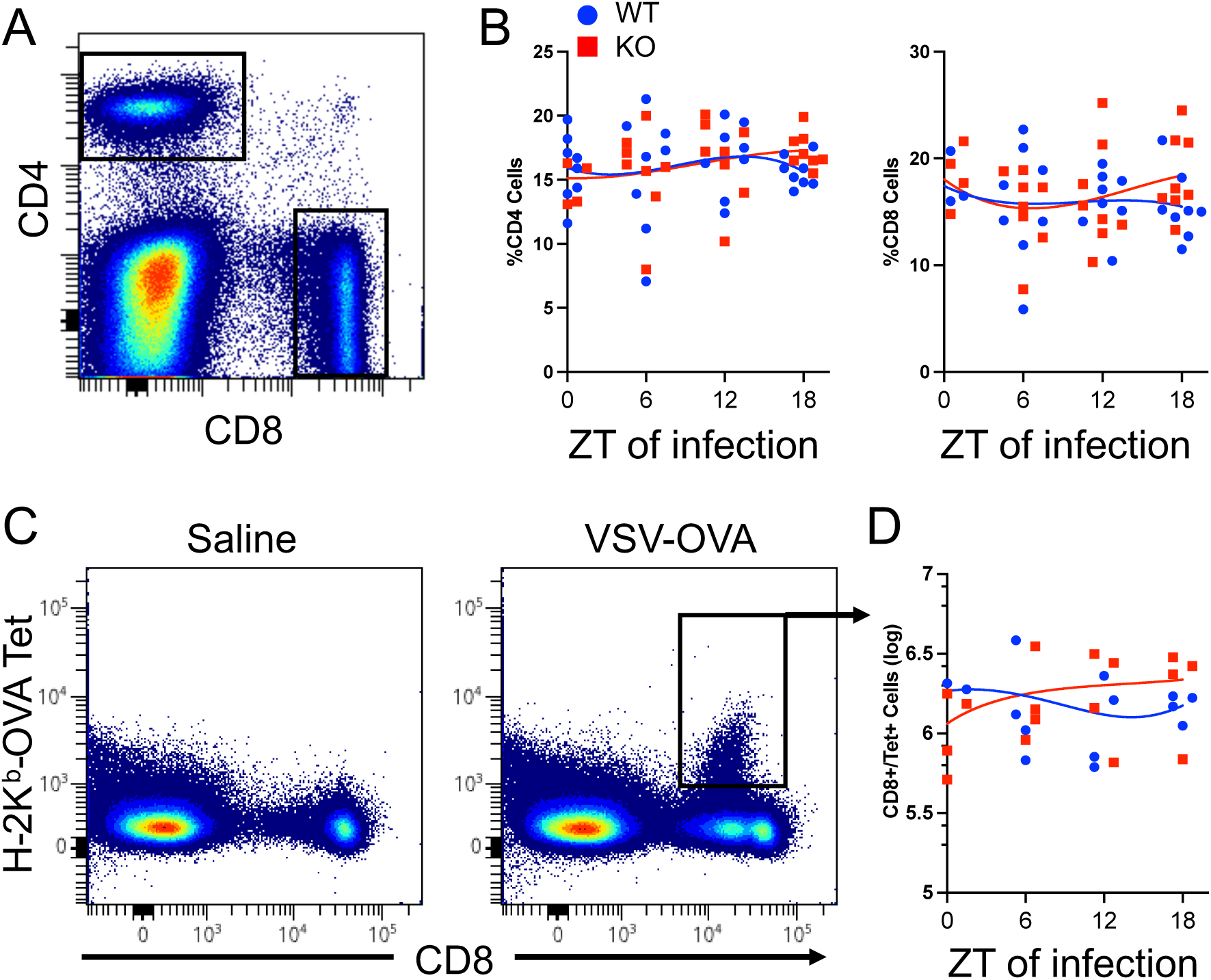
Expansion of Ag-specific CD8^+^ T cells as a function of time. WT and *Ad-fx-Cre* cohorts (n=4/group) were acclimated to 12:12 L/D for 2 weeks. Mice were infected with VSV-OVA at the indicated times and allowed to recover to d7. (A) Spleen cells were stained for CD4 and CD8 and analyzed by FACS. (B) The percent of CD4 and CD8+ splenic T cells were quantified from each cohort and displayed as a function of time of day of infection. (C) Representative dot plots of splenocytes stained for CD8 and K^b^-OVA tetramer are shown for animal infected with VSV-OVA, or with saline. (D) The absolute numbers of splenic K^b^-OVA-Tet+ cells were quantified and displayed as a function of time of day of infection. Circacompare analysis found no statistical rhythmicity in any of the populations described above (p > 0.05).

If the *Adrb*2 did not affect the numbers of Ag-specific responding cells, perhaps it controlled the quality of the cells as they developed into virus-specific effectors. To test this possibility, we stained cells with a panel of 8 additional antibodies specific for cell surface proteins that distinguish various subsets of cells based on their functional properties. We chose to include CCR7, CD62L, CXCR5, CXCR3, CD27, PD1, KLRG1, and CD44, as these markers identify numerous subsets of CD8^+^ T cells that fall in to the naïve, terminal effectors, effector memory, central memory phenotypes (29, 30). Batch analyses with tSNE and Phenograph of all samples identified 14 unique clusters based on marker expression (Fig. 7A), and the relative expression of each marker within the clusters is shown by heatmap analysis in Fig. 7B. These clusters represent unique cellular phenotypes within the responding antigen-specific population of CD8^+^ T cells, which are highly variegated based on their differential marker expression. This is expected based on previous work characterizing the profound temporal changes that occur in CD8^+^ T cells as they develop in response to virus infection (33). In assessing the abundance of each cluster, we find that some clusters (e.g. cluster 1) were rhythmic in their developmental response to virus, and their rhythmicity was similar between WT and Ad-fx-Cre (KO) cells (Fig. 7C). In contrast, other clusters were differentially rhythmic between WT and KO cells. For example, cluster 2 was rhythmic in WT but not in KO cells (Fig. 7D). Each of these clusters are distinguished by their differential expression of specified cell surface markers (Figs. 7C-E, right panels). Not only are these clusters heterogeneous in their phenotype, they are heterogeneous in their developmental regulation by the *Adrb2* and CRs. These data demonstrate that the differentiation of unique phenotypes of cells responding to virus at specific time points are regulated by intrinsic expression of the *Adrb*2 on CD8^+^ T cells.

**Figure 7.**
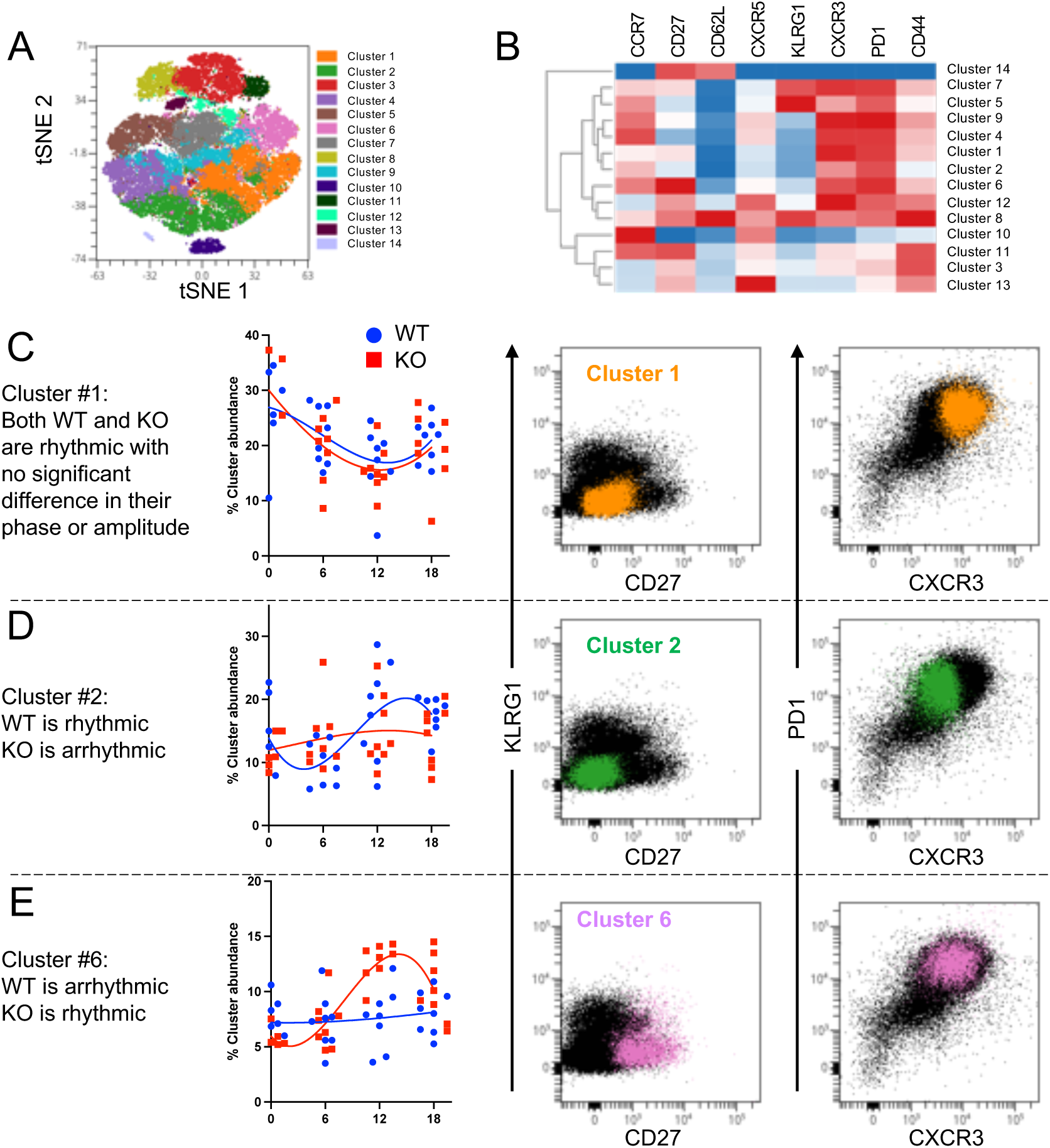
The *Adrb2* regulates circadian-dependent T cell development to virus infection. Splenocytes from mice in the experiment shown in Fig. 4A were stained with K^b^-OVA tetramer and a panel of 8 cell surface markers listed in paned B (n=8 animals/genotype/ZT of infection). (A) Cells were gated on live/CD8^+^/Kb-OVA^+^ cells, and all samples were batch processed through Phenograph, which identified 14 unique clusters. (B) Heat map of marker-level expression of clusters identified in panel A. (C-E) The percentage of clusters #1, #2, and #6 at each ZT of infection are displayed along with the rhythmicity results comparing WT to KO cells. The dot plots on the right are overlays of each cluster on top of the total parent population in order to indicate the relative expression of specific cell surface markers that define their phenotype.

All of these experiments to this point utilized VSV as a model virus, which is convenient but not particularly relevant to common human infections. For this reason, we have turned to influenza to determine whether *Adrb2*-driven CR regulation of T cell responses could be revealed broadly in other viral infections. Similar to the VSV infection model described above, we infected animals with IFA (H1N1, PR8) at different times of day, every 6 hours beginning at ZT0. All animals were sacrificed on d7 at the same time of day, and spleen and bronchial lymph nodes were harvested for analysis (Fig. 8). We assessed the unique T cell phenotypes by flow cytometry as in Fig. 7 and included a flu-specific tetramer, H-2D^b^-PA, which identifies CD8^+^ T cells reactive to the polymerase epitope of IFA. Again, we observed a robust expansion of flu-specific CD8^+^ T cells in the spleens of infected animals, however neither the percentages nor the total numbers of tetramer-reactive cells differed as a function of time of day of infection.

**Figure 8.**
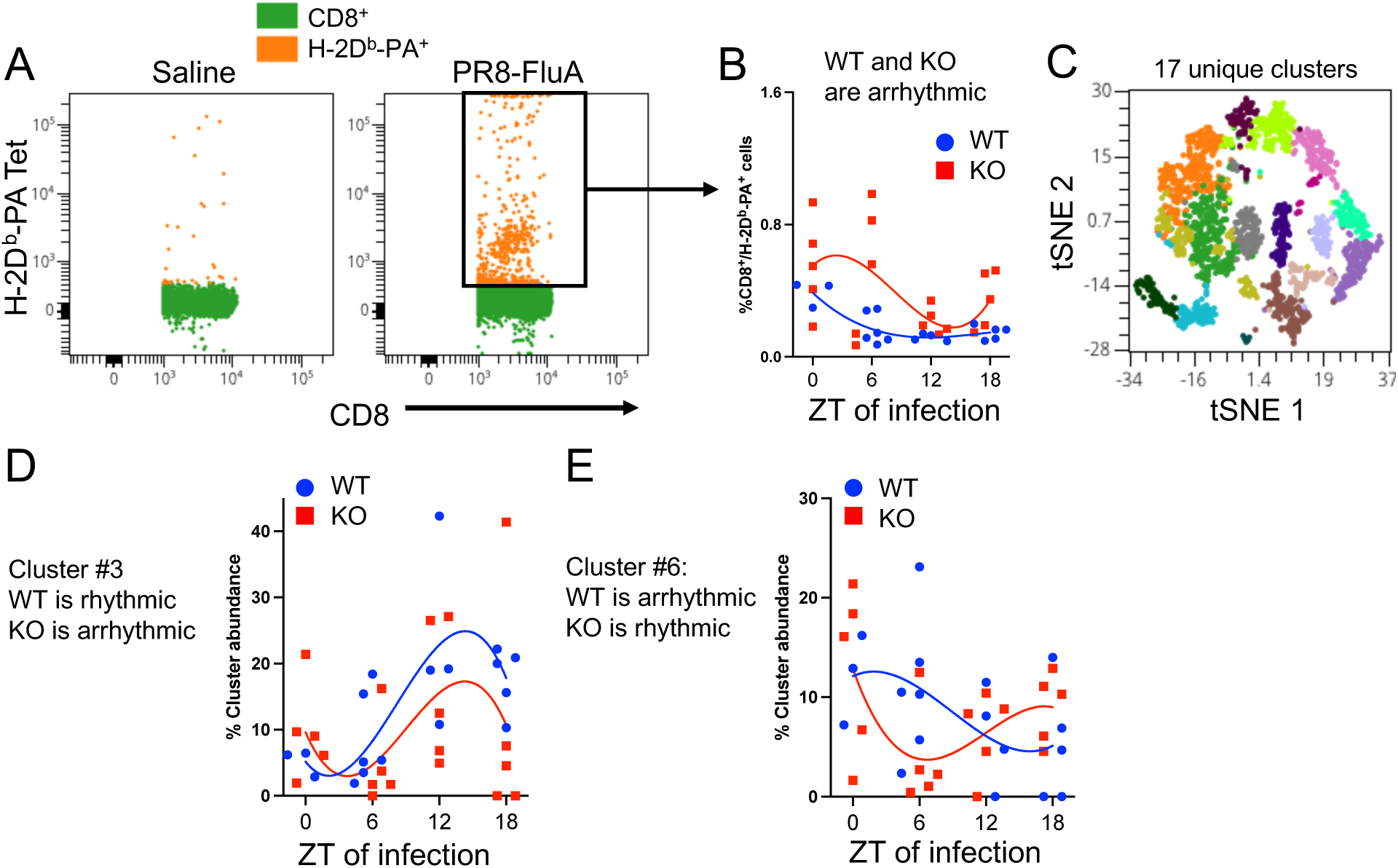
The *Adrb*2 regulates circadian-dependent development of flu-specific CD8^+^ T cells. WT and *Ad-fx-Cre* cohorts (n=4-5/genotype/ZT of infection) were acclimated to 12:12 L/D for 2 weeks. Mice were infected with IFA (PR8 strain) at the indicated times and allowed to recover to d7. Spleen cells were stained with α-CD8 + D^b^-PA-Tetramer along with 8 additional cell surface Abs described in Fig. 7 and analyzed by FACS. (A) D^b^-PA-Tet+ cells were gated and the percentage of antigen-specific cells were quantified as a function of ZT of infection (B). (C) Db-PA-Tet+ cells were clustered with Phenograph based on expression of cell surface marker, and cluster relatedness distributed on tSNE plots. (D-E) Relative abundance of clusters #3 (D) and #6 (E) were plotted, and rhythmicity as a function of ZT of infection was determined by Circacompare.

Cluster analysis of Tet+ cells identified 17 unique subsets (Fig. 7C) based on differential expression of the markers listed in Fig. 7. As we observed with VSV, IFA drove the differentiation of some clusters that expanded rhythmically as a function of time of day of infection. For example, cluster 1 displayed rhythmic expansion in response to IFA, and ablation of the *Adrb*2 in CD8^+^ T cells disrupted rhythmicity in this cluster (Fig. 8D). In contrast, cluster 6 was arrhythmic in WT cells but adopted a unique rhythm of expansion in KO cells (Fig. 8E). Thus, the *Adrb2* regulates the time-of-day-dependent balance of T cell phenotypes that emerge in response to IFA infection, and this mode of regulation is a novel mechanism to drive T cell developmental variegation.

## 3. Discussion

In this study, we report two major findings that link adrenergic signaling to entrainment of oscillations in CD8+ T cell gene expression and responses to viral infection. First, we found that intrinsic expression of the *Adrb2* is required for normal timely CR-dependent gene expression. Both innate and adaptive immune cells display rhythmic behavior such as homeostatic recirculation through lymph nodes and time-of-day-dependent responses to pathogens (4, 34–38). Most studies of CR regulation of immune function in mice have focused on innate immune cell responses to endotoxin (34, 39–44). However, recent studies have focused on CR regulation of adaptive T and B cells (19, 20, 45) and have even linked the *Adrb2* to this regulation (7, 46, 47). In mice, trafficking of B and T cells into lymph nodes follows a circadian oscillation, with both B and T cells optimally immigrating into lymph nodes toward the end of the light cycle (37). Immigration to LNs is regulated by diurnal oscillations in glucocorticoids, norepinephrine, and deletion of the *Adrb2* partially blocks the oscillatory nature of recirculation (7, 46). In some circumstances, oscillatory behavior in one cell type is directed by reciprocal oscillations in others. For example, circadian oscillation within dendritic cells regulates rhythmic responses to vaccine antigen in T cells by modulating antigen processing (48). A similar indirect mechanism of CR-dependent antitumor response by CD8+ T cells has also been observed (49). T cells also display direct CR-dependent behavior (21), which has been revealed by disrupting the core clock by genetic ablation of *Bmal1* (19, 20). Although various neurotransmitters such as glucocorticoids and norepinephrine have been shown to regulate these CR-dependent immune processes, their mechanism of control remained elusive.

Soluble neurotransmitters play a key role in regulating the light/dark entrainment of CR-dependent biological functions. In studying the role of epinephrine and norepinephrine in immune cell regulation, we found a remarkable immunosuppressive function for NE through ADRB2 signaling that potently inhibited the inflammatory processes of both CD8^+^ T cells (13) as well as innate macrophages and dendritic cells (14). In the course of these studies, we discovered an unexpected role for the *Adrb2* in directly regulating the CR core gene transcriptional pathway (22), and a detailed analysis of this finding is presented in the current study. We further show that intrinsic expression of the *Adrb2* on CD8^+^ T cells regulates the diurnal oscillation of many core CR genes and downstream CR gene targets. This is the first demonstration of a direct signaling pathway for the entrainment of CR gene oscillation in immune cells.

*Adrb2*-mediated CR gene regulation was selective, as not all CR genes were affected by the absence of ADRB2 signaling such as *Per1*. Although oscillation of the core CR genes *Per2* and *3* was disrupted, we observed several genes, which became oscillatory in expression or adopted a unique phase or amplitude in the absence of the *Adrb2*. There are several interpretations of this data. First, the curious acquisition of new frequencies by other genes could be explained by the continued response to other entrainment neurotransmitter receptors, such as the glucocorticoid receptor (GR) and the acetylcholine receptor (AChR), which are still intact and available for signaling in *Adrb2*-fx cells. Thus, in the absence of the *Adrb2*, continued oscillatory signaling through other entrainment receptors may artificially promote alternate expression patterns in downstream CR genes. Alternatively, there may be an intrinsic dysregulation that occurs when the normal oscillation frequency of one factor becomes disturbed in the absence of the *Adrb2*. For example, *Fbxl3* levels dropped dramatically at ZT+10 in knock-out cells. Given the modest rise in PER2 protein levels at ZT +10 in knock-out cells without a concomitant increase in mRNA, it is possible that the reduced levels of *Fblx3* at that time point created an artificial rise in PER2 protein. Similarly, a direct alteration in Per oscillation in the absence of *Adrb2* may indirectly impact *Bmal*, as these two factors reciprocally regulate each other. Finally, it is also likely that post-translational mechanisms, including phosphorylation and ubiquitination, are also under the direct control of Adrb2 signaling, which may impact both mRNA and protein levels of select CR genes. In conclusion, we find that the *Adrb2* is critical for regulating the oscillation of CR genes in CD8^+^ T cells under homeostasis.

Our second major finding demonstrates a clear role for the *Adrb2* in regulating CR-dependent diversification of T cell subsets that develop in response to virus infection. We found a remarkable time-of-day-dependent development of unique T cell subsets that displayed lineage marker expression consistent with populations of central memory T cells, which were dysregulated in the absence of the *Adrb2*. In WT mice, these CD27-/KLRG1-/PD1hi/CXCR3int cells peaked in their expansion at ZT12-ZT18, while their rhythmic development was lost in the absence of the *Adrb2*. The differential rhythmic nature we see in cellular phenotypes between WT and KO cells mirrors the types of rhythmic patterns we observed in gene expression. Some genes were rhythmic in WT but not in KO, and visa versa for others. It is likely that the loss of the *Adrb*2 allows other entrainment pathways to dominate and create alternative rhythms for some genes and for some developmental cell types. Recent studies have shown that IFA infection follows a clear CR-dependent inflammatory response in the lungs, with more severe inflammation occurring at later ZTs of infection (50, 51). Due to the critical involvement of CD8^+^ T cells in general responses to viral infection and IFA in particular, we propose that CD8^+^ T cell regulation by the *Adrb2* will control CR-dependent responses and inflammation. Our previous studies described a potent role for the *Adrb2* in acutely suppressing CD8^+^ T cell cytokine secretion and lytic activity (13). It is possible that some of the immunosuppressive activity of *Adrb2* signaling are independent of its regulation of CR gene oscillations, as both systemic and local levels of NE follow a diurnal pattern of secretion by sympathetic neurons that innervate secondary lymphoid tissues. *Bmal1* and *Per2* represent the core factors on either side of the clock pendulum, both of which are dysregulated in their periodicity in the absence of the *Adrb2*. Since whole-body deletion of *Bmal1* leads to enhance flu-mediated lung inflammation (52), we predict that *Bmal1*, and perhaps *Per2*, may also regulate the magnitude and timing of memory T cell responses to influenza in a manner that is dependent on entrainment by ADRB2 signaling. The current study opens new areas to explore the mechanisms underlying time-of-day dependent responses to infection and vaccination. Future studies will identify specific CR gene-dependent pathways in CD8^+^ T cells that regulate both their early effector cell development and later memory cell diversification and responses to recall infection.

## 4. Materials and methods

### Animal care

All mice used in this study were handled according to the guidelines of the Institutional Animal Care and Use Committee at the University of Texas Southwestern Medical Center. Wild type C57Bl/6J and the CD8α-Cre transgenic mice (27) were purchased from The Jacksons Laboratory. *Adrb2^fx/fx^*animals were provided by Dr. Gerard Karsenty from Columbia University (26). *Adrb2^fx/fx^* and Cd8αCre^+^ mice were bred to obtain Adrb2*^fx/fx^* Cd8αCre+ mice, and Cre-littermates were used for controls. The PER2::LUC+/+ mice (catalog number 006852) were received from Dr. Joseph Takahashi (28). The Adrb2*^fx/fx^* and Cd8αCre+ and PER2::LUC+/+ was bred to obtain PER2::LUC+/-Adrb2*^fx/fx^* Cd8αCre+ mice. All mice were housed and bred in specific pathogen-free facilities under a standard 12:12hr light/dark cycle with ad libitum feeding.

### Circadian Cabinets

All circadian studies were performed in a BSL2-level biohazard containment facility equipped with circadian cabinets. For both steady-state measurements and timed infections, animals were co-housed in cages placed in light/dark cabinets (Actimetrics). Each cabinet was programmed to a 12:12hr light/dark cycle differing by 6 hrs with respect to ZT0, initiating the light phase.

### Infections

For standard VSV infections WT and Adrb2 ^fx/fx^ Cd8αCre+ mice were infected by I.P. injection with 100 μl of either 10^6^ pfu of VSV-OVA (53) or saline. For influenza infection, mice were lightly anesthetized with isoflurane, and 30 μl of saline or 100 pfu IFA (A/PR8) was instilled into the nares. Animals were monitored and weighed daily until termination of the experiment.

### Luciferase assay

CD8 T cells were isolated from spleens of mice using CD8+ T cell purification kit according to the manufacturer’s instructions (Miltenyi Biotech). Luciferase activity was measured in purified CD8 T cell lysates with the luciferase assay system (Promega). Purified CD8 cells were lysed using cell lysis buffer 1X, and 100 ul of luciferase assay substrate was added to 20 ul of cell lysate in luminometer tubes. The tubes were vortexed briefly and luciferase activity was measure with a TD 20/20 luminometer (Turners) with the 2-second measurement delay followed by a 10-second measurement read for luciferase activity.

### RT-PCR analyses

Total RNA was isolated from CD8+ T cells with Qaigen RNA extraction kit, and 40-100 ng of RNA was used to perform reverse transcription using ABI High Capacity cDNA Reverse Transcription Kit (Applied Biosystems, Foster City, CA). The cDNA was used as template for the qPCR reactions using Bio-Rad’s SYBR® Green supermix using QuantStudio 7 Flex Real-Time PCR System (Applied Biosystems, Foster City, CA). The primers used to quantify mRNA are listed in Table 1. *Rn18s* and *Eif2a* were used as reference gene in qPCRs, and the average change in both references were used to calculate relative specific mRNA differences between samples. The relative gene expression differences were calculated using the 2^-ΔΔCt^ method (54).

### Flow and mass cytometry

Single cell suspensions of mouse splenocytes were used for staining in all flow cytometry measurements. For standard cell surface phenotyping, splenocytes were first stained with MHC-I tetramer (K^b^-OVA or ^Db^-PA) in RPMI media for 30 min at room temperature. The cells were then stained with a cocktail of cell surface antibodies (Biolegend, Table 2) in 0.5%BSA/PBS and analyzed on a LSR II flow cytometer (BD Biosciences). Data were compensated on FlowJo software followed by dimensionality reduction by OptSNE and clustering with Phenograph within the OMIQ data analysis pipeline (GraphPad).

**Table 2:** The list of antibodies used in staining of cells.

| Antibody | Fluorochrome |
| --- | --- |
| CD27 | BV421 |
| CD62L | BV510 |
| CD8 | BV570 |
| CXCR5 | BV650 |
| CD4 | BV711 |
| KLRG1 | FITC |
| CD44 | PerCP |
| CXCR3 | PE |
| PD1 | PE-Cy7 |
| TCR (CD8) | APC |
| CCR7 | AlxF700 |

For intracellular cytokine measurements by mass cytometry, splenocytes were first stimulated with PMA (50ng/ml) and Ionomycin (1mM) for 4 hours, with the addition of monensin for the last 2 hours of stimulation. Cells were then stained with APC-conjugated Kb-OVA tetramer at 37^0^C for 1 hour followed by staining for cell surface proteins with metal-conjugated antibodies listed in Table 3. Cells were then fixed and permeabilized with Fix-perm buffer (Standard Biotools) and stained for intracellular protein, i.e., cytokines. Cells were analyzed on a Helios mass cytometer (Standard Biotools), and data were normalized and downstream analysis performed on the OMIQ data analysis pipeline (Graphpad).

**Table 3:** The list of antibodies used in Mass Cytometry.

| Antibody | Metal |
| --- | --- |
| CD185 (CXCR5) | 142 Nd |
| CD69 | 143Nd |
| CD4 | 145 Nd |
| CD8 | 146 Nd |
| CD11b | 148 Nd |
| CD19 | 149 Sm |
| CD27 | 150Nd |
| CD25 | 151 Eu |
| CD3 | 152 Sm |
| TCRg | 159 Tb |
| CD62L | 160 Gd |
| anti-APC | 163 Dy |
| CD197/CCR7 | 164 Dy |
| Ki67 | 168 Er |
| TCRb | 169 Tm |
| NK1.1 | 170 Er |
| CD44 | 171 Yb |
| CD127/IL7Ra | 175 Lu |
| B220 | 176 Yb |
| CD11c | 209 Bi |
| TNFα | 141 Pr |
| IL2 | 144Nd |
| IL10 | 158 Gd |
| IFNγ | 165 Ho |
| IL6 | 167 Er |
| Perforin | 172 Yb |
| IL17A | 174 Yb |
| GranzymeB | 198 Pt |

### Statistical Analysis

Statistical tests were performed using GraphPad Prism version 9 software. One-way or two-way ANOVA was used followed by a Bonferroni posttest for pairwise comparisons within the groups. A Student’s two-tailed t-test was used for simple pairwise comparisons, differences with p>0.05 were considered significant. The rhythmic pattern of oscillation and the differences in mesor, phase and amplitude were quantified by CircaCompare (55) statistical tool available on Github.

For high dimensional flow and mass cytometry data, all populations were either identified by Boolean gating (cytokine-secreting cells) or by clustering (cell surface phenotypes). For clustering, data were first subjected to dimensionality reduction by OptSNE and then clustered with the Phenograph algorhythm at k=50. SAM tools of OMIQ were used to perform statistical comparisons between groups and between time zones. Data from statistically significant comparisons were then analyzed for rhythmicity with Circacompare.

## Acknowledgments

The authors wish to thank Angela Mobley and the UT Southwestern Flow Cytometry core for help with flow and mass cytometry. We thank Dr. Lora Hooper for excellent suggestions on experimental design and help with the BSL2 circadian laboratory.

This work was supported by NIH grants AI175217, AI143248, and AI125545.

