## Supplemental Information for "The β2-adrenergic receptor (ADRB2) entrains circadian gene oscillation and diurnal responses to virus infection in CD8^+^ T cells"

### Supplementary Materials:

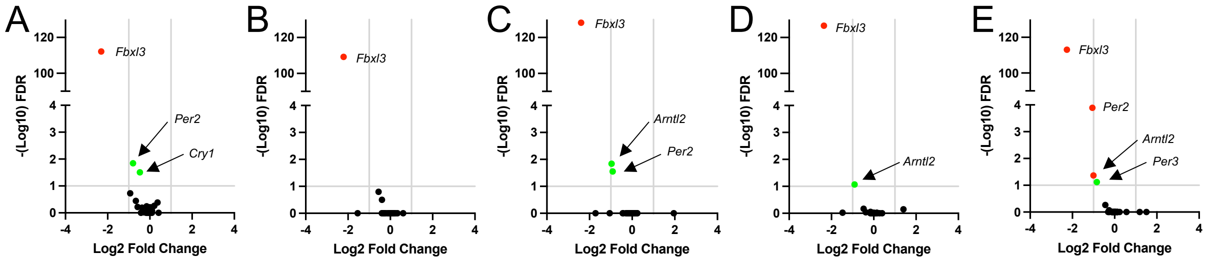


**Supplemental Figure 1. Differential core clock gene expression in *Adrb2*$^{-/-}$ CD8$^{+}$ T cells in response to virus infection.** Volcano plots representing differentially expressed core clock genes described in the heat map in Fig. 1A. Each graph displays gene expression in CD8$^{+}$ T cells isolated on d0, d4, d5, d7, and d12 (panels A-E respectively) after VSV-HA infection in Estrada, et. al., (22). Green symbols are significantly different at <2-fold compared to WT cells; red symbols are significantly different at >2-fold compared to WT cells.

| Rhythmicity | Genes | Shape *p* value | Rhythmicity *p* value |
|---|---|---|---|
| **WT Arrhythmic**<br>**KO Arrhythmic** | Clock<br>Cry1<br>Tpl2 | - | >0.05 |
| **WT Arrhythmic (p>0.05)**<br><br>**KO Rhythmic (see rhythmicity p value)** | Nr1D1<br>Nr1D2<br>Nfil3<br>NfKB<br>Rora<br>Otd3<br>GranzymeB<br>IFN AR1<br>Perforin<br>Nfat2 | - | 1.38E-03<br>7.81E-05<br>5.03E-03<br>1.22E-02<br>1.43E-02<br>2.07E-02<br>3.27E-02<br>1.42E-02<br>4.36E-02<br>3.22E-02 |
| **WT Rhythmic (see rhythmicity p value)**<br><br>**KO Arrhythmic** | Per2<br>Per3<br>Cry2<br>Bmal2<br>EomesTbr2<br>TbetTbx21<br>TNF<br>IL12Rb2 | | 1.34E-02<br>9.87E-03<br>6.52E-03<br>4.52E-02<br>1.43E-02<br>1.23E-02<br>2.96E-03<br>2.25E-02 |
| **WT & KO Rhythmic with significant difference in phase, mesor, or amplitude** | Bmal1<br><br>Fbxl3<br><br>Nr4a1 | phase 6.99E-02<br>mesor 3.58E-02<br>amplitude 2.99E-01<br><br>phase 4.62E-02<br>mesor 3.22E-01<br>amplitude 7.65E-01<br><br>phase 9.57E-02<br>mesor 2.63E-02<br>amplitude 2.38E-01 | WT, 1.12E-02; KO, 3.66E-04<br><br>WT, 7.52E-04; KO, 3.37E-02<br><br>WT, 4.98E-02; KO, 2.49E-04 |
| **WT & KO Rhythmic without significant difference** | Per1<br><br>IFN-g | phase 1.9E-01<br>mesor 7.4E-01<br>amplitude 0.3E-01<br><br>phase 6.31E-01<br>mesor 6.56E-01<br>amplitude 2.78E-01 | WT, 1.01E-03;<br>KO, 1.433E-03<br><br>WT, 1.89E-03;<br>KO, 1.17E-02 |

**Supplemental Table 1**. **Statistical analysis of gene expression rhythmicity in purified CD8+ T cells.** Data displayed in Figs. 2 and 3 were assessed for rhythmicity and aspects of wave form including phase, mesor, and amplitude by the Circacompare statistical package. Genes are categorized based on their differences in rhythmicity between WT and Adrb2$^{-/-}$ cells.
